# *Staphylococcus aureus* virulence factor expression matters: input from targeted proteomics shows Panton-Valentine leucocidin impact on mortality

**DOI:** 10.1101/2022.09.18.508069

**Authors:** Mariane Pivard, Sylvere Bastien, Iulia Macavei, Nicolas Mouton, Jean-Philippe Rasigade, Florence Couzon, Romain Carrière, Karen Moreau, Jérôme Lemoine, Francois Vandenesch

## Abstract

In the case of commensal bacteria such as Staphylococcus aureus, the transition from commensalism to invasion and disease as well as disease severity in the course of an infection remain poorly predictable on the sole basis of virulence gene content. To determine whether variations in the levels of expression of the numerous *S. aureus* virulence factors could affect disease occurrence and/or severity, we developed a targeted proteomic approach that monitored 149 peptide surrogates targeting 44 proteins. Semi-quantification was achieved by normalization on the signal of ribosomal proteins. We then evaluated this approach on a series of *S.aureus* strains from 136 patients presenting a severe community-acquired pneumonia, all admitted to an intensive care unit. After adjusting to the Charlson Comorbidity Index score the multivariate analysis of severity parameters found that HlgB, Nuc, and Tsst-1 were positively associated while BlaI and HlgC were negatively associated with leucopenia. BlaZ and HlgB were positively associated with hemoptysis and HlgC was negatively associated with hemoptysis. Regarding mortality, both the multivariate (1.28; 95%CI[1.02;1.60]) and survival (1.15; 95%CI[1.016;1.302]) analyses showed that only PVL was associated with death in a dose-dependent manner. Beyond highlighting the decisive role of PVL in community-acquired pneumonia severity, this study brings the proof of concept that “expression matters” and proposes a method that can be routinely implemented in laboratories, for any Staphylococcal disease, and which could be developed for other commensal bacteria.

**One Sentence Summary:** A highly multiplexed semi-quantitative mass spectrometry method was developed for 44 *Staphylococcus aureus* virulence factors; applied to a 136-strain collection from severe community-acquired pneumonia patients, it showed that Panton-Valentine leucocidin was the only factor to impact mortality in a dose-dependent manner.

## INTRODUCTION

*Staphylococcus aureus (S. aureus)* is detected in 30% of the population, mostly as a nasal commensal, but also in the throat, on the skin, and in the gastrointestinal tract [1]. In addition to this innocuous interaction with the human host, *S. aureus* has the potential to develop a wide range of diseases in humans, from mild infections of the skin and soft tissues to severe and fatal infections, such as bacteremia and pneumonia [2]. The transition from the commensal status to the pathogenic one is attributed to both host and environment parameters and to the capacity of *S. aureus* to produce a wide diversity of virulence factors such as adhesins, toxins, innate immune evasion proteins, and other effector proteins. However, whilst all *S. aureus* strains encode a core set of major virulence factors, such as protein A, coagulase, and Phenol Soluble Modulins (PSMs) [3–5], the virulence gene content is not enough to predict the transition from commensalism to invasion and disease, nor can it predict disease severity. Moreover, although certain accessory genome-encoded virulence factors are specifically associated with certain diseases, such as superantigenic toxins (e.g. TSST-1, SEA,..) with toxic shock syndrome [6,7], or exfoliative toxins with bullous impetigo and scalded skin syndrome [8], the *S. aureus* strains containing these specific genes can also colonize the host without inducing any disease. Overall, this suggests the expression level of these factors must also be considered. A recent study showed that assessing the cytotoxicity level and biofilm-forming abilities *in vitro* on THP1 cells can help predict mortality in *S. aureus* bacteremia in a lineage-dependent manner. Elevated cytolytic toxicity combined with low levels of biofilm formation was predictive of an increased risk of mortality in infections by strains of the CC22 but not CC30 lineage [9]. Another study showed that strains from pneumonia (community- or hospital-acquired) were more cytotoxic for neutrophils than nasal-colonizing strains [10]. A study assessing the *in vitro* level of two regulatory RNA (RNAIII and sprD) identified a significant but marginal correlation between these markers and severity in the course of S.aureus bacteremia [11]. Finally, high *in vitro* expression of alpha-hemolysin (Hla) was associated with ventilator-associated pneumonia (VAP) whereas low Hla producing strains were associated with ventilator-associated tracheobronchitis [12]. Overall, there have been very few attempts to determine whether variations in the expression level of the numerous *S. aureus* virulence factors could affect disease occurrence and/or severity in the course of an infection. High-throughput approaches to RNA and protein quantification are limited by cost-effectiveness and human resource constraints, impeding researchers from studying the correlation between the expression levels of a large panel of virulence factors and the severity and mortality data obtained from large clinical cohorts.

To overcome these issues, we developed a targeted proteomic approach, a technology of choice for implementing highly multiplexed protein assay methods. The strategy consists in selecting a list of peptide surrogates issued from the enzymatic digestion of bacterial proteins according to so-called top-down proteomics. Thus, an assay method targeting 44 virulence factors and resistance markers was deployed on a triple quadrupole mass spectrometer operating in Multiple Reaction Monitoring (MRM). After technical validation of the method, the approach was evaluated on a series of *S. aureus* isolates from 136 patients presenting a severe community-acquired pneumonia (CAP), all admitted to an intensive care unit. The results of this study show a decisive role of PVL in community-acquired pneumonia severity and brings the proof of concept that the level of expression must be taken into account when considering Staphylococcal pathogenesis.

## RESULTS

### MRM assay development and choice of the peptide surrogates

A list of 44 virulence factors and resistance markers, including the most commonly accepted candidates [4,13], was designed (Table S1). The first phase of the assay development aimed at designing the reporter peptide candidates to be tested with respect to their detectability in a validation set of 20 *S. aureus* strains. All predicted proteotypic sequences obtained by *in silico* trypsin digestion and containing between 5 and 25 amino acids were retained as candidate peptides. Although methionine or tryptophan residues have a propensity to undergo uncontrolled oxidations that may impair the sensitivity of the assay method [14], peptides containing these residues were nevertheless retained to expand the panel of peptides sampled. Conversely, peptides containing cysteine residues were systematically excluded to avoid the reduction and alkylation steps of the disulfide bridges, which could complicate the sample preparation protocol in the perspective of being able to implement this assay method as a routine test in a near future.

MRM assays targeting the panel of peptide candidates were built using the Skyline web tool without *a priori* knowledge of the chromatographic retention times (https://skyline.ms/project/home/software/Skyline/begin.view). As usually recommended, only transitions corresponding to Y type fragment ions were retained to build the testing methods, owing to their propensity to induce the most intense and less interfered signals. No more than 50 peptide candidates, i.e. 150 transitions, were included in each MRM assay so as to ensure that a minimum of 8 data points were collected per reconstructed ion chromatogram. A manual inspection of the raw data using the Skyline tool enabled to establish an intermediate list of peptide reporter candidates. A peptide reporter was considered putatively detected when the retention times of its respective transitions were perfectly aligned on the chromatogram and when it was detected in at least one strain of the validation set. Among the 44 virulence factors, at least one peptide reporter candidate was pinpointed for 33 proteins. In the specific case of PSMα1 and PSMα4 polypeptides, the short sequences and the location of arginine and lysine residues imposed the selection of only one and two peptides, respectively, that were therefore directly synthetized. For the remaining 9 proteins (Table S1, red characters), no peak could be confidently assigned on the chromatogram, indicating that these virulence factors were either not expressed or expressed at a concentration below the detection limit in the strains of the validation set. Given their potential relevance in pathogenesis, these proteins were maintained in the final assay. Hence, the synthetic forms of all peptides, were then monitored in order to accurately validate their retention time and optimize the ionization and fragmentation parameters. Since the absolute retention time of each peptide was accurately assessed, it enabled the building of a unique scheduled MRM method targeting the 149 reporter peptides (Table S1).

### The proteome of clinical strains is highly diverse

The MRM assay was applied to a series of 136 isolates collected from a French cohort of patients with community-acquired pneumonia (Table S2) [15]. Since all the strains had been sequenced, they were all initially checked to ensure that the peptide sequences targeted in the final method covered all potential allelic variations for each protein. The results of the targeted proteomics revealed a broad variation in the expression of each of these factors ranging from 2 log2 (e.g. SEA) to almost 8 log2 for numerous proteins such as Spa, SdrD, Nuclease, Hld, PVL, and others (Figure 1). However, for 7 proteins (EdinC, ETA, ETB, MecC, SEB, SED, and SEH), only very few strains produced the proteins or the levels were too low to allow detection. These proteins were *a posteriori* removed from the analysis to avoid unnecessary background noise due to the lack of variance between strains. As the quantifier of Spa targets the repeat region of this protein, relative quantification may be biased depending on the number of repeats in the strains. However, no correlation was found between Spa expression and the number of repeats in each strain (Figure 2). In addition, the differences of up to 8 log2 (i.e. 256 fold) observed between strains are much higher than the 1 to 3 fold expected differences based on the number of repeats. Therefore, Spa quantification was retained in the analysis without controlling for the number of Spa repeats in the quantification values.

**Figure 1.**
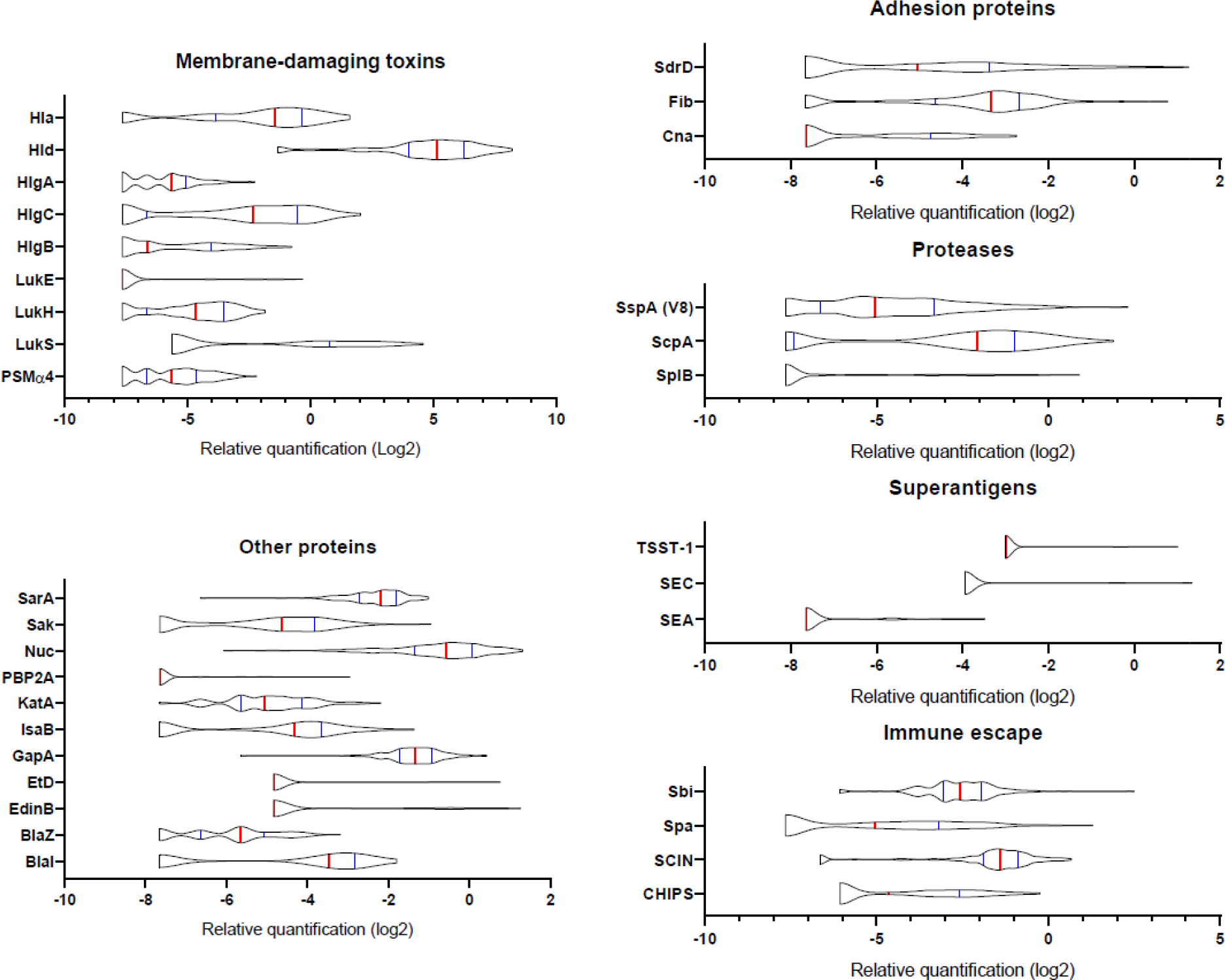
Variations in the expression of virulence factors between clinical strains. The relative amount of virulence factors expressed in log2 and categorized into functional groups are represented using violin plots. A total of 136 clinical strains were analyzed. The red vertical lines represent the medians and the blue lines represent the 1^st^ and 3^rd^ quartiles.

**Figure 2.**
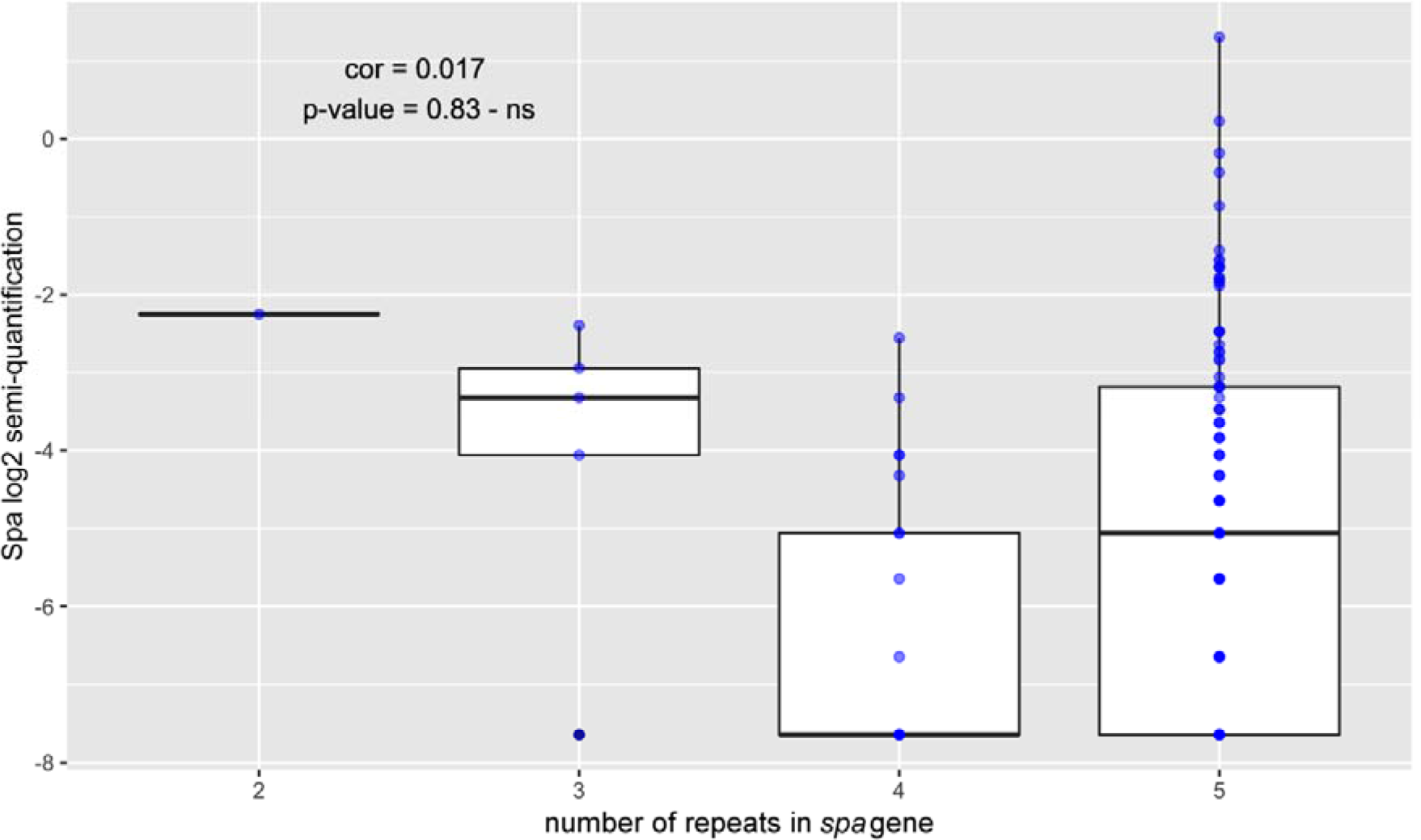
The number of repeats in the *spa* gene does not impact its quantification. The relative amount of log2-transformed Spa is represented by dot plots (blue dots) according to the number of immunoglobulin-binding repeats present in the spa gene of each strain, determined using genome sequencing. Medians with 25^th^ and 75^th^ percentiles are represented by box plots. A Pearson correlation test between Spa semi-quantification and the number of repeats was performed, p-value > 0.05 - ns.

Given the large diversity in the genetic background of the corresponding strains ([15] and Table S2), the possibility that the proteome could be correlated with the genetic background was tested by principal component analysis. Neither the four *accessory gene regulator (agr)* types nor the 21 clonal complexes (CC) and 2 sequence types (ST) of the collection explained the proteome distribution as no proteomic profile could be associated with any of these genetic backgrounds (Figure 3). We then searched for correlations between the 37 proteins by constructing a correlation heatmap (Figure 4). Strong correlations were observed for numerous virulence factors reflecting the fact that they are encoded in operons. This was the case for LukS and LukF (+0.98) of the PVL operon, LukE and LukD (+0.726) of the LukDE operon, and PSMa1 and PSMa4 (+0.780) of the PSM operon. The absence of correlation between HlgC and HlgB (+0.240) of the HlgCB operon however, likely reflects post-transcriptional regulatory steps on this operon. Strong correlations were also observed for proteins whose expression are immediately dependent on a common regulatory pathway, as is the case for the PSMs and Hld which are the first targets downstream of the agr activation pathway. Conversely some unexpected correlations were observed such as that of EdinB with Exfoliative toxin D (+0.72), LukG with SplB (+0.610), and Spa with SspA (V8 protease) (−0.59).

**Figure 3.**
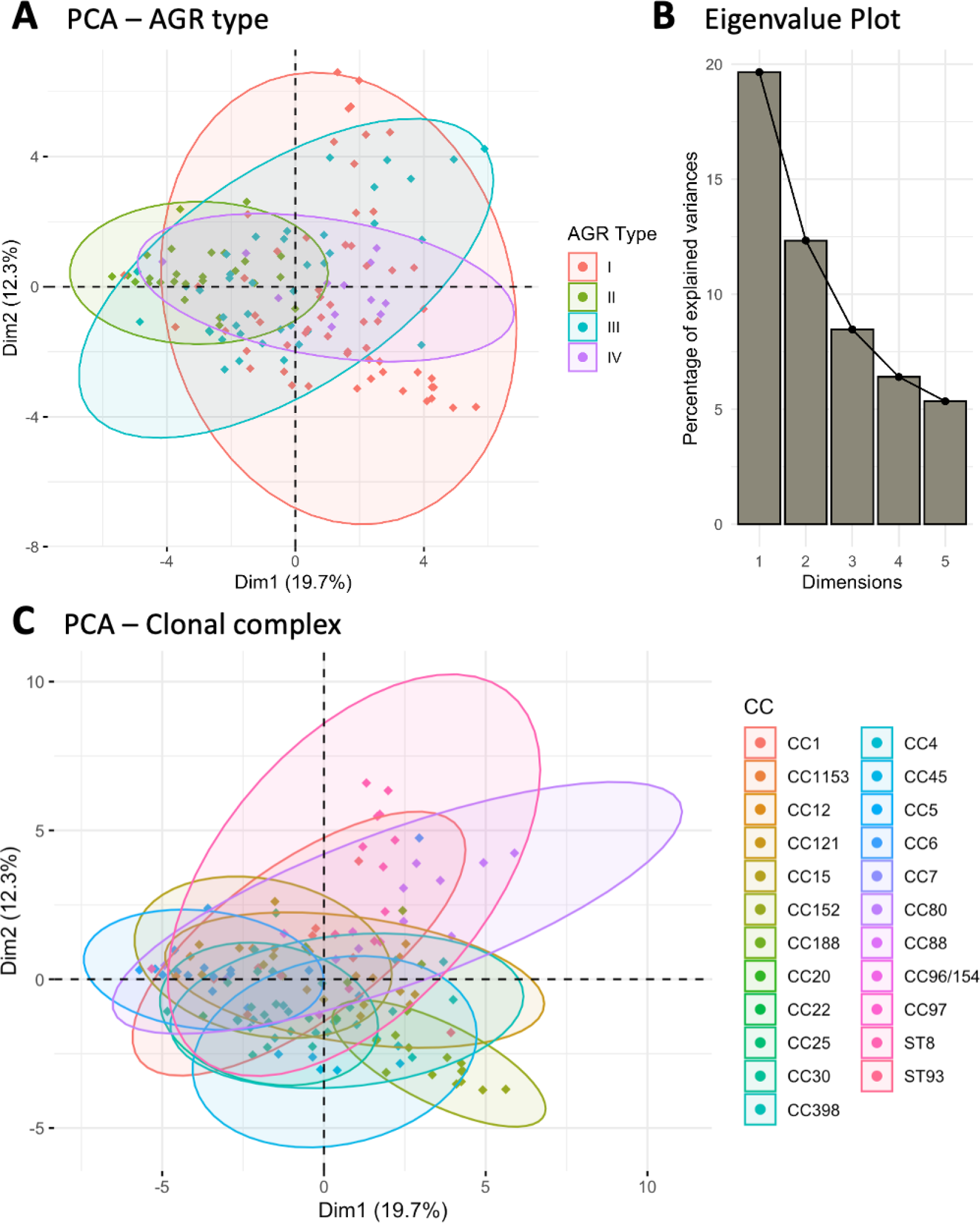
*S. aureus* proteome is not linked to the strains’ genetic backgrounds. Principal component analysis (PCA) on the log2-transformed relative protein quantities was conducted. (**A**) The variations in the virulomes depending on the 1^st^ and 2^nd^ dimensions of the PCA analysis are visualized according to the AGR type. (**B**) The associated Eigenvalue plot for the first five dimensions of the PCA is represented using a histogram plot. (**C**) The variations in the virulomes depending on the 1^st^ and 2^nd^ dimensions of the PCA analysis are visualized according to the clonal complex (CC).

**Figure 4.**
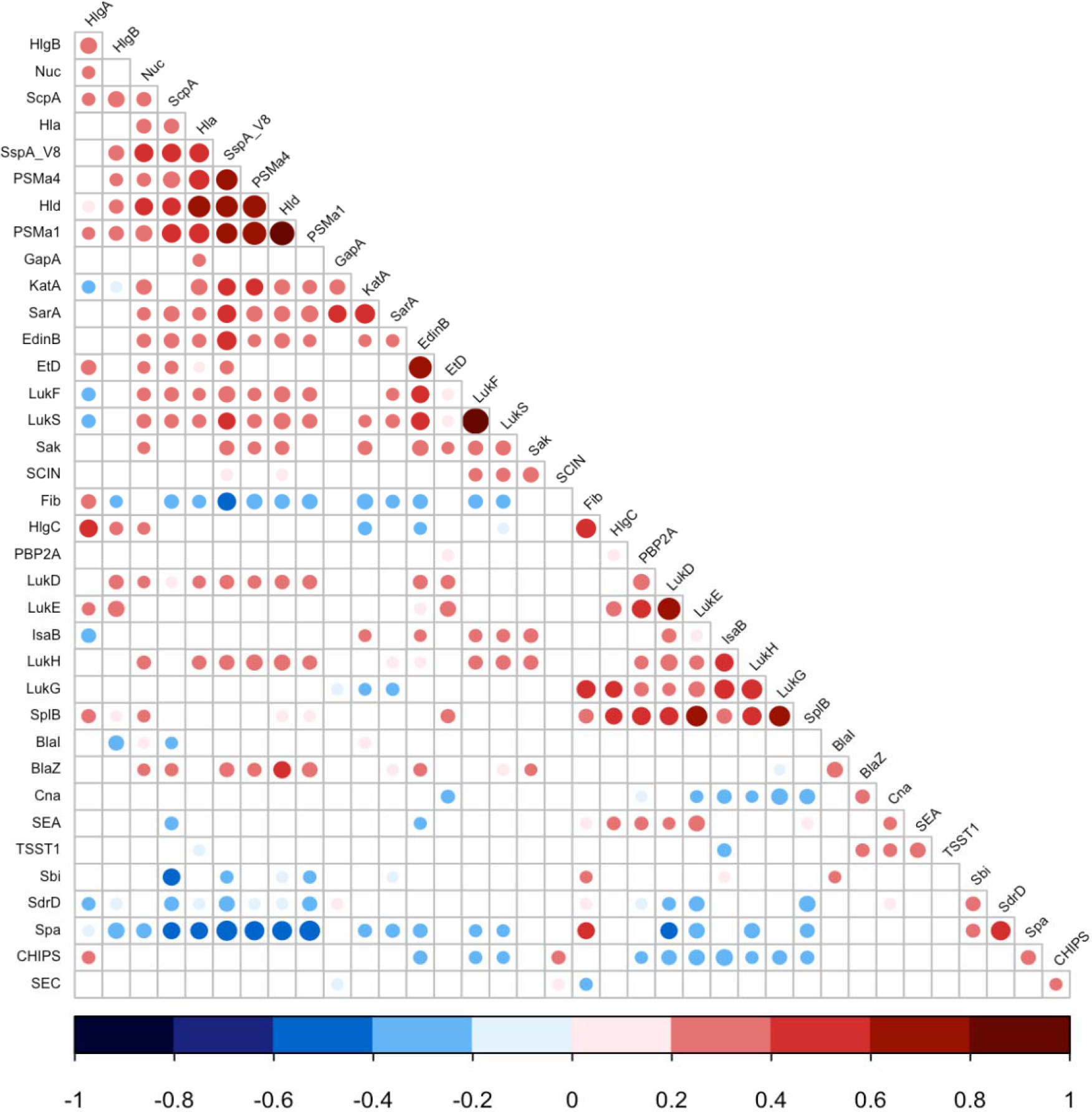
Expression correlations between *S. aureus* virulence factors. Linear correlations between the expression of 37 virulence factors were tested using Pearson correlation test. Non-significant correlations are represented as empty squares and significant correlations as dots. The dot size is negatively proportional to the p-value. The Pearson coefficient values are represented as dark blue for negative values to dark red for positive ones.

### PVL impacts mortality in a dose-dependent manner

The previous study of strains from the current patient cohort aimed to decipher the relationship between the presence/absence of genes encoding virulence factors and CAP outcome. The study showed that PVL was an independent predictor of death, hemoptysis, and leukopenia. It also found that *edinB* (encoding the epidermal differentiation inhibitor) was a predictor of hemoptysis and leukopenia and that the staphylococcal enterotoxin A (seA) gene predicted leukopenia [15]. We thus examined the possible association of the same severity parameters (death, hemoptysis, and leukopenia) with the proteomic data, assuming that such an analysis based on quantitative data could reveal novel associations, particularly regarding the virulence factors present in all strains. To increase the robustness of the analysis, we first took into account the multi-collinearity of several factors, notably those encoded in operons, except when they were not strongly correlated as was the case for HlgC and HlgB. The final list of factors was tested in univariate and multivariate regression, controlled on the Charlson score to account for age and co-morbidities. The univariate analysis highlighted several virulence factors associated with severity (positively or negatively) in a dose-dependent manner. Those that were associated with death were PVL, Pbp2A, Glutamyl endopeptidase (V8 protease), BlaZ, Hla, PSMα1-4, EdinB, LukH, Hld, SdrD, and Fib (Figure 5A). Those associated with hemoptysis were EdinB, PVL, BlaZ, Hld, Glutamyl endopeptidase (V8 protease), Nuc, EtD, LukH, and Spa (Figure S1A), and those associated with leukopenia were Glutamyl endopeptidase (V8 protease), EdinB, Nuc, Hld, PVL, HlgB, SspP (staphopain), PSMα1-4, Etd, Sak, SarA, Hla, LukGH, SCIN, Spa, and Fib (Figure S2A). However, in multivariate regression analysis only PVL remained associated with death in a dose-dependent manner (odds ratio 1.28 95% CI 1.02 to 1.60; Figure 5B). HlgB, Nuc, and Tsst-1 were positively associated with leucopenia while BlaI and HlgC were negatively associated with leucopenia (Figure S2B). Finally, BlaZ and HlgB were positively associated with hemoptysis and HlgC was negatively associated with hemoptysis (Figure S1B). When restricting the analysis to PVL-positive strains/patients, PVL production was significantly higher in deceased versus non-deceased patients (Figure 6). In multivariate survival analysis, only PVL was associated with death (1.15 95% IC 1.016 to 1.302; Table S3). To elucidate the fact that many potentially important virulence factors (e.g. Hla) were neutralized in the multivariate analysis, we looked for a possible correlation between the expression of these factors and that of PVL for each strain. The expression of Hla and of several factors identified in univariate analysis was significantly correlated with the expression of PVL in PVL-positive strains (Figure 7 and S3 and S4), thus demonstrating that these factors were actually confounding variables.

**Figure 5.**
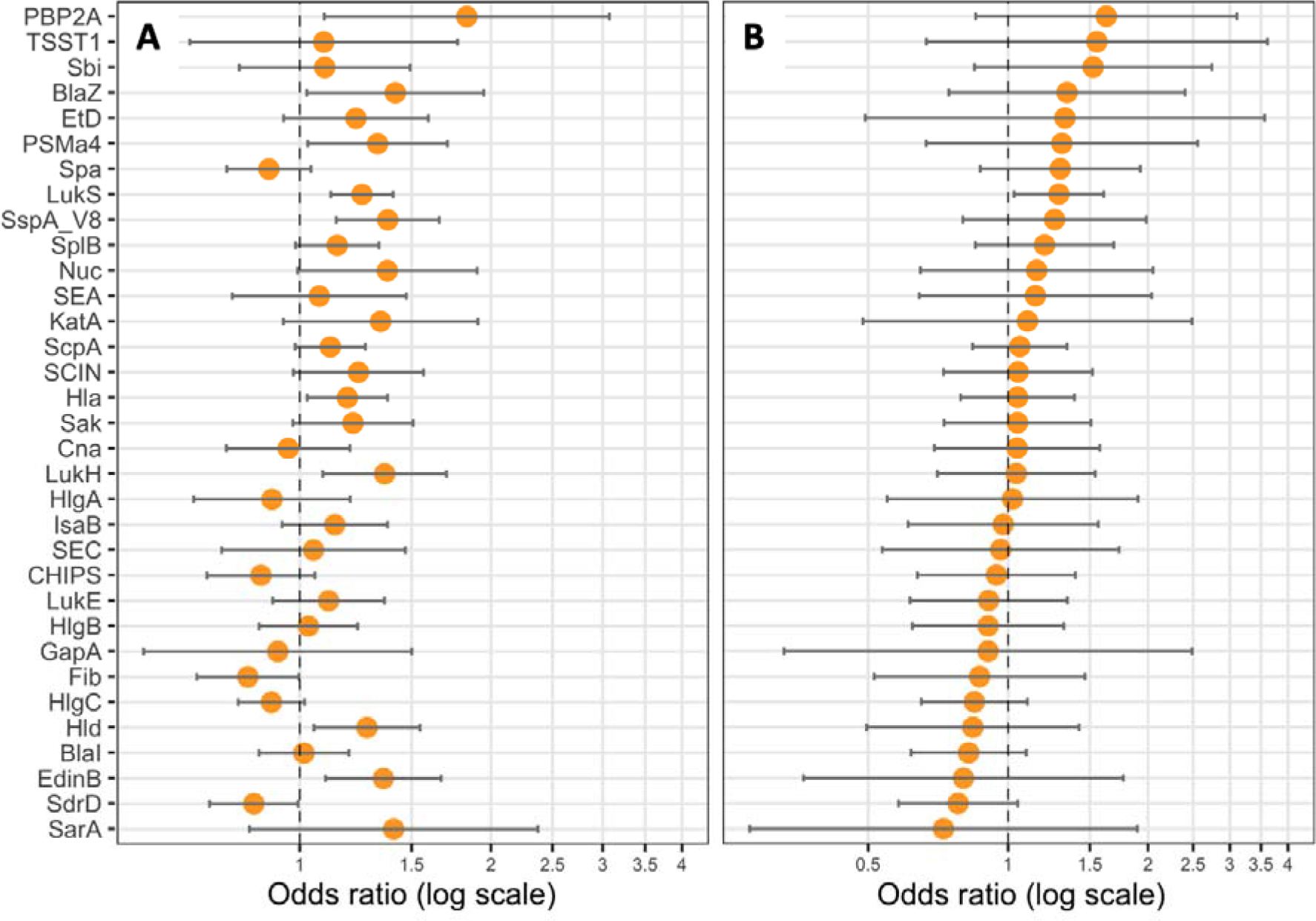
PVL (LukS) expression is correlated with death. Logistic regression models including Charlson score as covariate were performed, with either the expression level of (**A**) one protein (univariate), or (**B**) all proteins (multivariate) as explicative variables, and death as the response variable. Coefficients were reported as odds ratios (orange dots) with 95% confident intervals (error bars).

**Figure 6.**
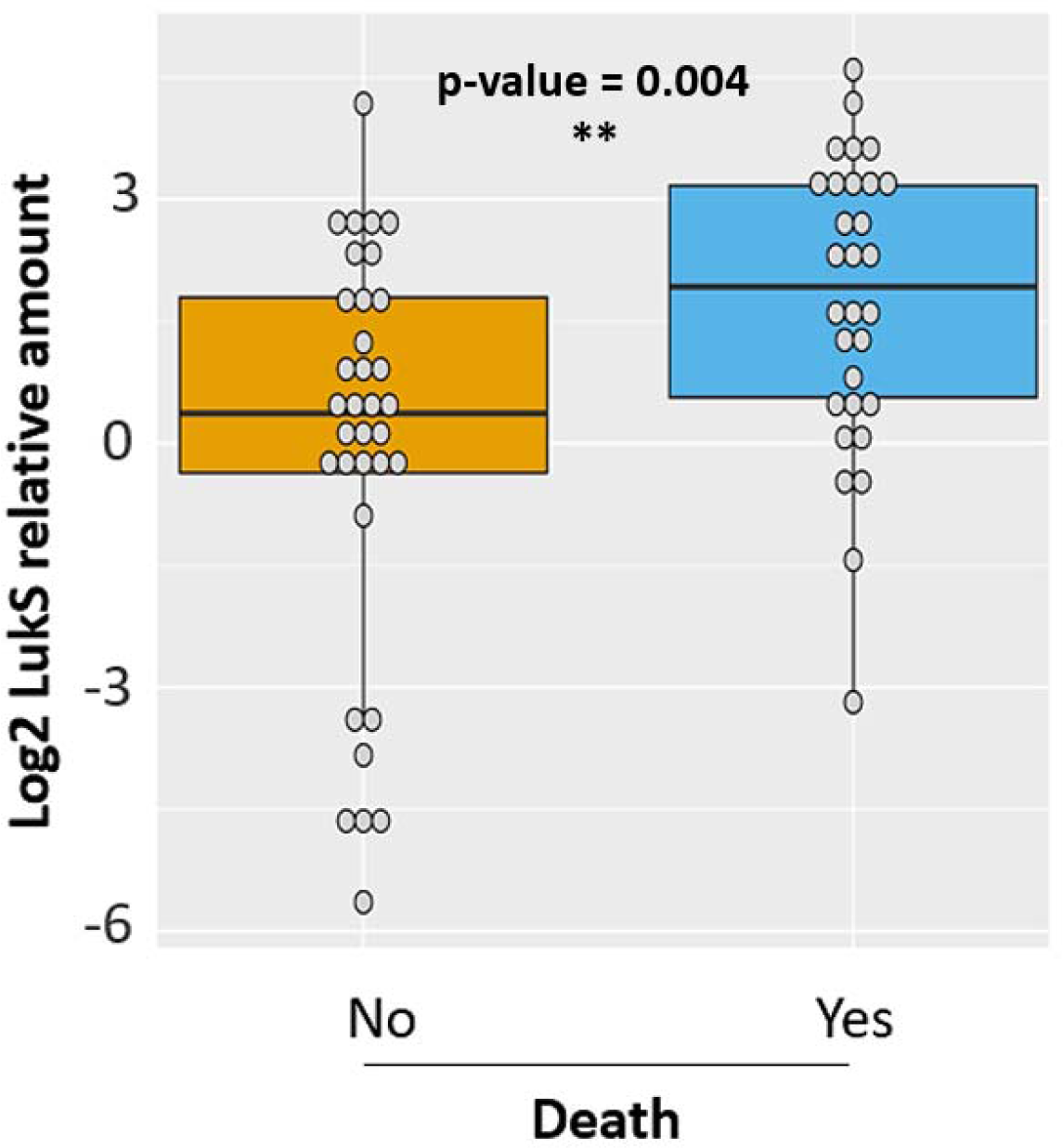
Strains isolated from deceased patients produce more PVL. Relative amount of log2-transformed LukS from PVL-positive strains isolated from deceased (blue, n=30) and no-deceased (orange, n=34) patients are represented by dot plots. The medians with 25^th^ and 75^th^ percentiles are represented by box plots. A logistic regression model including CCI as covariate was performed, p-value < 0.01 - **.

**Figure 7.**
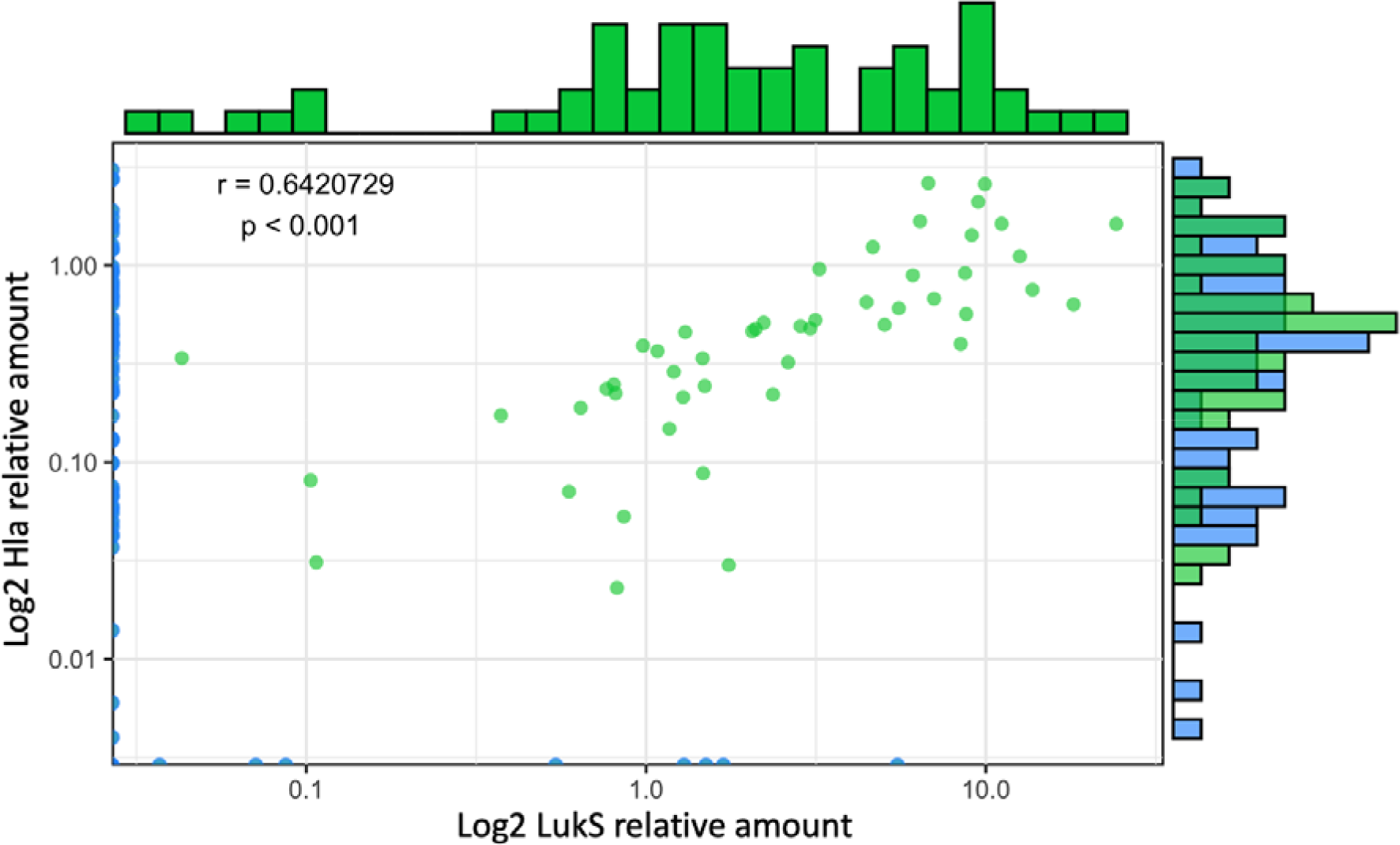
The expression levels of Hla and PVL are correlated. Log2-transformed Hla relative quantities were plotted against the log2-transformed relative quantities of LukS. The density of strains expressing similar protein quantities is represented by histograms, on top for LukS and on the right side for Hla, except for null values. Green dots represent strains with quantification for both Hla and LukS, and blue dots represent strains with at least one null value. A Pearson correlation test was performed.

### Functional clustering of virulence factors

*S. aureus* virulence factors are often functionally synergistic if not redundant without necessarily targeting the same receptors. For example, all *S. aureus* enterotoxins behave as superantigens that polyclonally activate T cells although each of these toxins targets a specific set of Vbeta T cell receptors. Similarly, PVL and *Hla* are pore-forming toxins that target different receptors but their pathogenic effect may be synergistic as evidenced by the lower mortality of rabbits infected with an hla mutant and infused with PVL immunoglobulins [16]. To test this hypothesis, 20/37 toxins were functionally clustered into five categories corresponding to adhesion proteins, membrane-damaging toxins, superantigenic toxins, immunoglobulin-binding proteins, and proteases (Table S4). These five categories were tested for their potential association with the studied severity parameters (i.e., death, hemoptysis, leukopenia) but no significant association was found (Table S4).

## DISCUSSION

In the present study, we developed a multiplexed scheduled-MRM assay targeting 162 peptides used as surrogates for the detection and quantification of a panel of virulence factors expressed by *S. aureus.* Since the objective was to evaluate a putative correlation between the expression level of these factors and the severity of human diseases, notably CAP, the repeatability and accuracy of the quantification process are critical to provide usable quantitative data. Although secondary, the objective of the methodological development stage was also to ensure that the assay technique could then be used routinely to characterize the strains sent to the French National Reference Center for Staphylococci and, in the future, to any clinical laboratory mastering mass spectrometry.

The vast majority of studies implementing targeted proteomic techniques for protein quantification describe the use of stable heavy isotope labeled (SIL) peptides as internal standards. Although relevant, particularly in the context of complex preparation methods as it compensates the bias due to the loss of sample [14], this approach has many limitations. The most obvious pertains to the cost of the labeled peptides if the method is highly multiplexed. The tedious longitudinal internal standard concentration quality controls, both in stock and working solutions, required to assess individual SIL peptide concentration, is another limitation worth mentioning. It should also be noted that the use of SIL peptides does not dispense with the need for normalization on protein quantity and thus on the number of cells processed. Considering the aforementioned specifications and drawbacks related to the use of SIL peptides, the normalization of the expression level of the virulence proteins was herein based on an internal standardization relative to ribosomal proteins. Indeed, in a context of non-limiting nutrient concentration, the ribosomal protein content is fine-tuned for fitness strategy allocation, and thus exhibits a linear relationship with growth rate and hence, cell number [17–19]. The relevance and reliability of this relative quantification strategy based on the absolute signal of peptides deriving from ribosomal proteins is supported by a previous study demonstrating a good correlation between the expression level of proteins involved in various antibiotic resistance mechanisms and the resistance phenotypes of bacterial strains [20].

The quest for robustness, repeatability, and affordability of an assay, which could eventually be implemented as a routine test, requires streamlining the number of steps of the pre-analytical sample processing which may be the source of bias. Compared to conventional protocols of bottom-up proteomics, the sample preparation was greatly simplified in the present approach. Indeed, the reduction and alkylation of the thiol group of the disulfide bond was omitted, cell lysis and trypsin digestion steps were performed concomitantly, and the time-consuming off-line or on-line peptide desalting prior to mass spectrometry analysis was avoided. Herein, this last step could be avoided because a robust micro-flow chromatographic format was preferred to the gold standard nano-chromatography encountered in most proteomics experiments. When the amount of starting biological sample is not limiting, the micro-flow format shows similar sensitivity while offering greater repeatability and robustness, especially when hyphenated with a triple quadrupole mass spectrometer. Such analytical platforms, which are already in place in certain hospitals for various clinical purposes, some being certified to perform In Vitro Diagnostics tests, could thus advantageously be used in the context of bacterial virulence assessment.

When applied to a large series of unrelated *S. aureus* isolates, the first noteworthy feature revealed by the approach was the diversity of protein expression levels, independently of strain lineage. This is consistent with a recent study showing significant lineage-independent variability in the cytoxicity of *S. aureus* strains [21]. Thus, the virulence of a given strain cannot be simply inferred from its virulence gene content and/or lineage. The coordinated expression of *S. aureus* virulence factors has been studied for decades, since the description and characterization of the major *agr* global regulon, followed by the discovery of a plethora of regulatory proteins and RNA (for review [22–25]). It is known that *agr* represses the expression of numerous surface proteins such as coagulase, ClfA, FnbpAB, Spa, and others, and upregulates the expression of several toxins including Hla, PVL, certain superantigens, nucleases, and proteases [26–29]. Hence, positive and negative correlations are expected within and between these two groups of factors, respectively, and were observed at the protein level in the present study. Moreover, several of the virulence factors studied herein are encoded in operons [30], which should lead to, if not an equimolar production, at least a strong positive correlation in the expression of the two proteins. This was not the case for HlgC and HlgB, suggesting that, depending on the strain, more complex regulation may occur at the level of transcription (e.g., *hlg*B transcription generated by an alternative promoter) or translation (e.g., HlgB or C translation efficiency may vary between strains due to subtle nucleotide variations in the RBS or upstream). We are currently investigating this hypothesis by use of genetic and molecular approaches.

CAP due to *S. aureus* is rare but severe and is often associated with the production of a rarely encountered toxin, PVL [31,32]. This toxin which targets the C5a receptor at the surface of granulocytes monocytes ,and macrophages [33] was shown to impact severity of *S. aureus pneumonia* in a murine model [34]. It was also found to be a risk factor for mortality in adolescent and adult patients with severe CAP, independently of methicillin resistance, *S. aureus* genetic background, and other virulence factors. However, it is important to note that mortality still remained important in patients infected with PVL-negative strains, who represented nearly half the cohort previously studied [15]. The contribution of core-genome encoded virulence factors can therefore not be dismissed, particularly since experimental studies have demonstrated the importance of Hla, Protein A, staphylococcal nuclease, and many other virulence factors in the pathogenesis of staphylococcal pneumonia (for review see [35]). The lack of dose-dependent correlation between these virulence factors and death observed herein might reflect the dominant effect of PVL and the limited size of the PVL-negative group. Regarding the case of HlgCB which targets the same receptor as PVL at the surface of myeloid cells [36,37], the former was anticipated to impact severity in a dose-dependent manner in PVL-negative strains. However, in multivariate analysis HlgB was positively associated with hemoptysis and leukopenia whilst HlgC was negatively associated with these markers of CAP severity. These apparently conflicting results may reflect the complexity of the interplay between these toxins or may be explained by the functional duality of HlgB which composes both the HlgAB and HlgCB toxins, each having specific biological functions [36]. However, it is worth mentioning that in patients infected with PVL-negative strains, mortality was 40% (6/15) in HlgC high level producing strains compared to 24% (14/57) in HlgC low level producing strains (data not shown). This suggests that the small sample size has likely prevented any significant association between HlgCB expression and death to be observed.

The association of nuclease with leucopenia is remarkable since nuclease has never been shown to have a direct impact on circulating neutrophils, even though it has been shown to degrade neutrophil extracellular traps, promoting immune cell death [38]. The association of TSST-1 with leucopenia observed herein is also novel in humans and could result from the extravasation of cells in a Vbeta-unrestricted manner, as demonstrated experimentally in rabbits [39]. The inverse association of BlaZ and BlaI with two severity parameters is in accordance with the molecular circuit of the *bla* operon since BlaI represses the transcription of *bla*Z encoding the Beta-lactamase BlaZ [40]. Beta-lactamase hyperproduction can reduce the efficacy of penicillinase-resistant penicillins commonly used for staphylococcal infections such as oxacillin and 1^st^ generation cephalosporins [41], thus causing treatment failure [42].

The lack of association with severity when grouping 20 virulence factors into five functional categories is unexpected, as the results of animal studies have shown an additive neutralizing effect of a therapeutic antibody targeting five pore-forming toxins [43]. Altogether, these results may reflect that CAP, as opposed to VAP in which Hla plays a major role [12], is a disease in which the main contributor to death is PVL. For this toxin, the correlation between the observed clinical severity and the *in vitro* expression level provides additional evidence for causality and not simple mere association. This causality was already suggested in rabbits using PVL isogenic mutants and therapeutic PVL-directed antibodies [16,34].

Although the 37 virulence factors studied herein include the most studied candidates, this relatively low number is a main limitation of the present study. This number was dictated by technical (multiplexing capacity) and economic constraints: the need to setup a cost-effective approach that can be applied to large sets of isolates. Moreover, we made the hypothesis that *in vitro* expression could predict an *in vivo* behavior. The fact that only the expression of PVL predicted death in the context of CAP legitimates this approach, at least for the virulence factors studied. However, the results herein cannot conclude firmly on the role of the virulence factors other than PVL, as the single *in vitro* condition used might not be optimal for all of them. Other culture media, possibly including various stress conditions, should be developed in future studies to improve the model. Another limitation is the choice of a cohort of patients with CAP to evaluate the potential value of this approach. We hope that the application of this method to other clinical conditions such as bacteremia, infective endocarditis, VAP, or bone and joint infections will uncover novel associations between disease presentation, complications, or outcome and the quantitative expression of virulence factors.

## MATERIALS AND METHODS

### Proteomic validation set

The 20 *S. aureus* strains were selected from the WG sequenced biobank of the French National Reference Center for Staphylococci. The rationale for strain selection was to obtain positive representative isolates for each of the virulence factors of interest, allowing to estimate the detectability of the method for each protein.

### Strains and data of the proteomic test set

Clinical isolates and accompanying clinical data were obtained from a previously published French cohort of severe CAP [15]. The clinical features retained for the present study were those representing severity and outcome in the cohort study: death, leukopenia (defined as total leukocyte count < 3 G/L), and hemoptysis extracted from the Gillet study [15] (Table S2). The age-adapted Charlson Comorbidity Index (CCI) score was retained as a control covariate to adjust prediction models to the baseline characteristics of the patients. Patients aged less than 3y (n=20), who were previously found to present very distinct clinical presentation and outcome compared to older patients, were excluded. Patients co-infected with several *S. aureus* strains (n=3) were also excluded. Finally, 4 isolates were excluded a posteriori because they harbored a rare allelic variant of the protein A which prevented its quantification by the detection method used (2 strains from deceased patients and 2 from patients who survived). Altogether, 136 isolates and patients were included in the final analyses (Table S2). All strains were whole genome-sequenced to validate the presence of virulence factor-encoding genes. DNA sequences are available at [Data not available yet on LENA].

#### Sample preparation

After thawing, each strain was grown in 4 mL of Casein Hydrolysate Yeast Extract (CCY) complemented with 10% pyruvate acid in 25 mL tubes at 37°C with agitation at 200 rpm. After 18h of growth, 50 μl of this culture was inoculated into a new CCY tube and grown for 8 hours. 1 mL of 8h-culture was collected in duplicate and stored at −80°C until a denaturation step at 90°C during 1h. Then, samples were stored at −80°C until mass spectrometry preparation and analysis. For both the analytical development phase and the strain collection analysis, series of 6 samples were prepared by pipetting 200 μL of inactivated bacterial cell suspension prepared into 1.5 mL Eppendorf tubes filled with approximately 70 μL of 150-212 μm glass beads (Sigma-Aldrich). 50 μL of a trypsin solution (Porcine pancreas grade, Sigma-Aldrich) extemporaneously prepared at a concentration of 1 mg/mL in 150 mM NH_4_HCO_3_ (Sigma-Aldrich) was added to digest the proteins released by sonication during 10 minutes applying irradiation cycles of 30 seconds at the low irradiation setting (Bioruptor ultrasonicator, Diagenode, Lièges Belgique). Importantly, the commercial version of the sonication device was modified to keep the bath temperature at 50°C instead of the conventional 0°C setting. Digestion was finally stopped by adding 5 μL of formic acid (Sigma-Aldrich) and the Eppendorf tubes were centrifuged at 10000 g during 5 minutes. 180 μL of the supernatant was finally transferred into 2 mL screw cap tubes (Labbox) endowed with a 250 μL glass insert.

#### Targeted mass spectrometry analysis

The Scheduled Multiple Reaction Monitoring (MRM) experiments were deployed on a hybrid triple quadripole/linear ion trap mass spectrometer 6500 QTRAP (AB Sciex, Toronto) equipped with an ESI Turbo V™ ion source operating at 550 °C and using an ion spray voltage of 5500 volts. The settings of curtain gas and nebulizing gas 1 and gas 2 were adjusted to 50, 70, and 60 psi, respectively. The instrument and the data acquisition and processing were controlled by the Analyst 1.6.2 software. Unit mass resolution was fixed for Q1 and Q3 in MRM mode.

The tandem mass spectrometer was hyphenated to a liquid chromatography set-up comprising 2 binary pumps (1290 series, Agilent technologies). For each experiment, 80 μL of the trypsin digest was injected and submitted to on-line Solid Phase Extraction on a PLRP-S, 12.5 mm x 2.1 mm column (Waters) connected to a Rheodyne 10-injection port valve. Peptides were desalted with 100% water containing 0.1% formic acid v/v (LC-MS grade, Fisher Scientific) for 3 minutes at a flow rate of 100 μL/min. After the valve switching, peptides were separated on a Waters BEH C18 column (100 mm x 2.1 mm; 3.5 μm particle size) using the following gradient: 98% water, solvent A (LC-MS grade, Fisher Scientific) containing 0.1% formic acid (v/v) and 2% acetonitrile, solvent B (LC-MS grade, Fisher Scientific) containing 0.1% formic acid (v/v) for 3 minutes followed by a 17-minute linear gradient to reach 35% of solvent B. The scheduling window was fixed at 1.2 minute around the expected retention time of the peptide targets, which kept the dwell time above 5 milliseconds and ensured at least 10 data points per reconstructed transition peak.

#### Peptide and protein detection validation

MultiQuant TM 2.1 software (AB Sciex) was used to process the raw data leading to automated chromatographic peak detection for each transition signal as well as the calculation of their absolute area (MQ4 algorithm). Peak detection parameters were set as follows: a Gaussian smooth width of 2.0 point, a retention time half window of 35 seconds, a baseline subtraction window of 2 minutes, and a 50% noise level for baseline. A reference transition ratio (TR) was calculated for each peptide from the synthetic form (equation 1), which validated effective detection of peptide surrogates during the analyses of the *S. aureus* strain collection. Only the peptides with a TR standard deviation below 20% relative to the mean TR were ultimately validated.

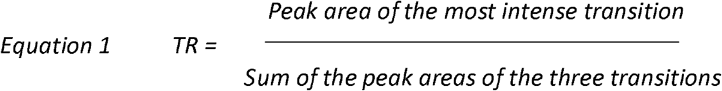

#### Protein quantification

It is assumed that the MS1 signal and the most intense transitions of the best flying peptides correlate with protein abundance [44] and therefore can be used as proxy for label-free quantification. To take into account any putative sensitivity deviation of the mass spectrometer instrument and variation of bacteria cell density in the suspension, quantification of each virulence protein was derived from the calculated ratio between the sum of area of the three transitions of a peptide quantifier of the virulence protein and the average of the sum of the three transitions of three peptides deriving from three ribosomal proteins (equation 2): IYPGENVGR (30S protein S11), EMSVLELNDLVK (50S protein L7/L12), ALQSAGLEVTAIR (50S protein L27). Ribosomal proteins were used as internal relative standard as their expression level correlates with cell number amplification during the exponential growing phase [17–19].

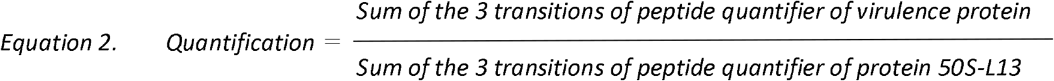

#### Statistical analysis

We used visualization and regression procedures to assess the relationships between the expression level of virulence factors, the isolate’s genetic background, and the clinical outcome of the infected patient. Relative protein quantities were log2-transformed before analysis to improve the interpretation of model coefficients, as this transformation allows to express coefficients as the change of the response variable per doubling of the protein quantity. To avoid obtaining infinitely negative values after transformation, relative protein quantities equal to zero were replaced with the smallest non-zero quantity for the same protein and divided by 2 before being log2 transformed. All analyses used R software v4.0.3 (R Core Team [2019]). The anonymized data and software code required to reproduce the results are available at [https://github.com/Mpvrd/Proteomic].

To visualize the variations of the virulome according to the strain genetic background, principal component analysis of the log2-transformed relative protein quantities was conducted. The linear correlation between the log2-transformed relative quantities of proteins was tested for significance using a Pearson correlation test.

The associations between clinical outcomes (death, leukopenia, hemoptysis) and the relative quantities of virulence factors were analyzed in logistic regression models. Univariate models were constructed separately for each protein, and multivariable models included all proteins at once to estimate their independent effect on the outcome. All models (including the separate univariate models) included the CCI as a covariate to adjust for patient health at baseline. To reduce the risk of model instability due to multicollinearity between protein quantities, strongly correlated proteins (Pearson correlation >0.5, p-value <0.05) encoded in the same operon were identified and only one protein of the operon, selected arbitrarily, was retained for statistical analysis. Coefficients of the logistic regression models were reported as odds ratios with 95% confidence intervals based on a normal approximation. The associations between relative protein quantities and the delay from hospital admission to death or hospital discharge were analyzed using Cox survival regression models, both univariate and multivariable, using the same predictors as the logistic regression models. Coefficients of the survival regression models were expressed as hazard ratios with 95% confidence intervals based on a normal approximation.

We examined whether groups of proteins, rather than individual proteins, collectively predicted clinical outcomes. To this aim, functionally-related proteins were grouped in categories listed in Table S4. For each of the previously described models, the point estimate of the pooled coefficients of proteins in a category was obtained as the sum of the coefficients of individual proteins. The variance of the pooled coefficients was obtained as the sum of the covariance matrix of the individual coefficients, as described elsewhere [45].

The variances of the pooled coefficients were then used to construct 95% confidence intervals based on the same normal approximation as the individual coefficients. These pooled coefficients were interpreted as the collective effect of the proteins in the category on the clinical outcome predicted by the model.

## List of Supplementary Materials

**Figure S1.**
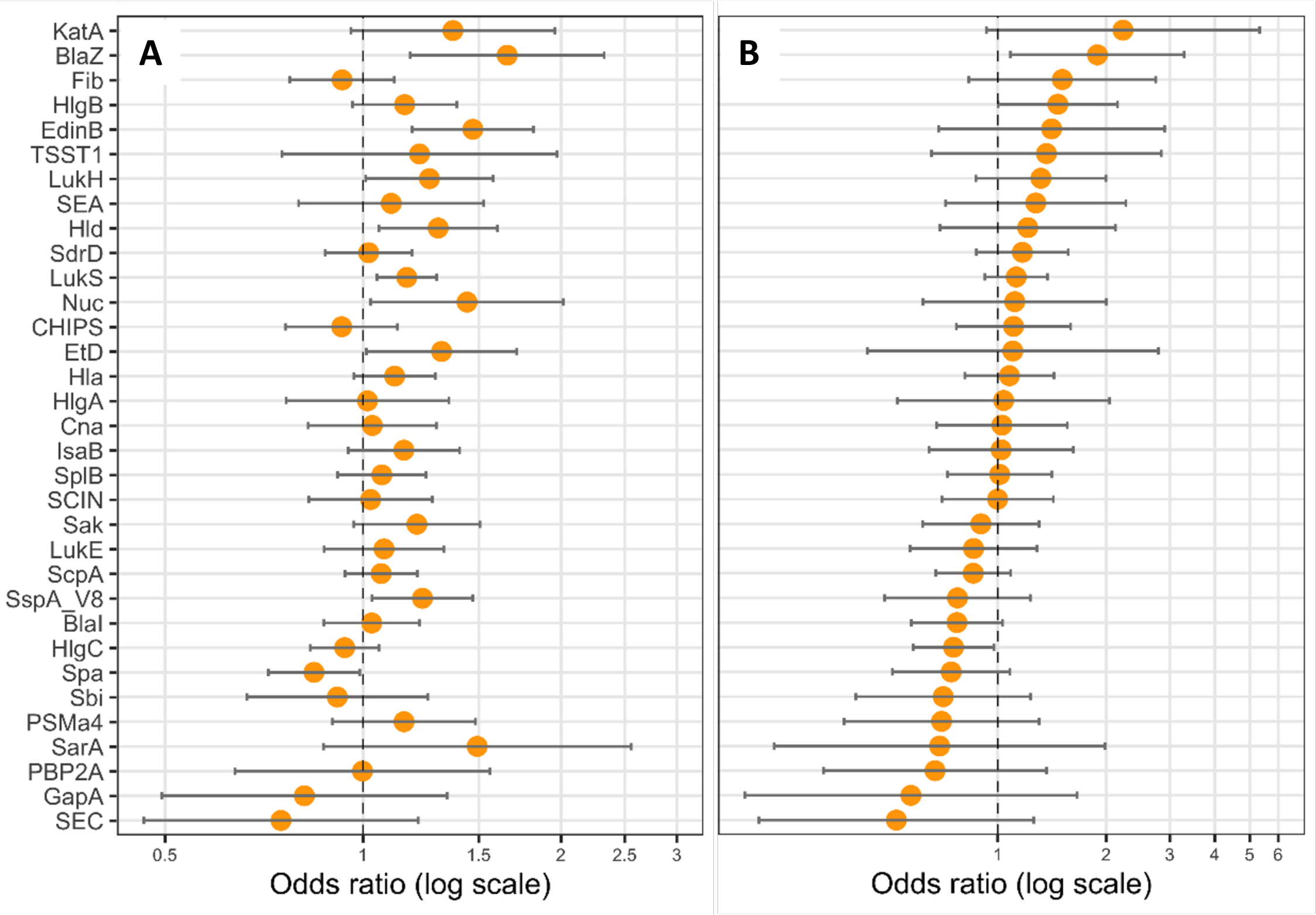
BlaZ and HlgB expressions are correlated with hemoptysis. Logistic regression models including CCI as covariate were performed, with either the expression level of **(A**) one protein (univariate), or (**B**) all proteins (multivariate) as explicative variables, and hemoptysis as the response variable. Coefficients were reported as odds ratios (orange dots) with 95% confident intervals (error bars).

**Figure S2.**
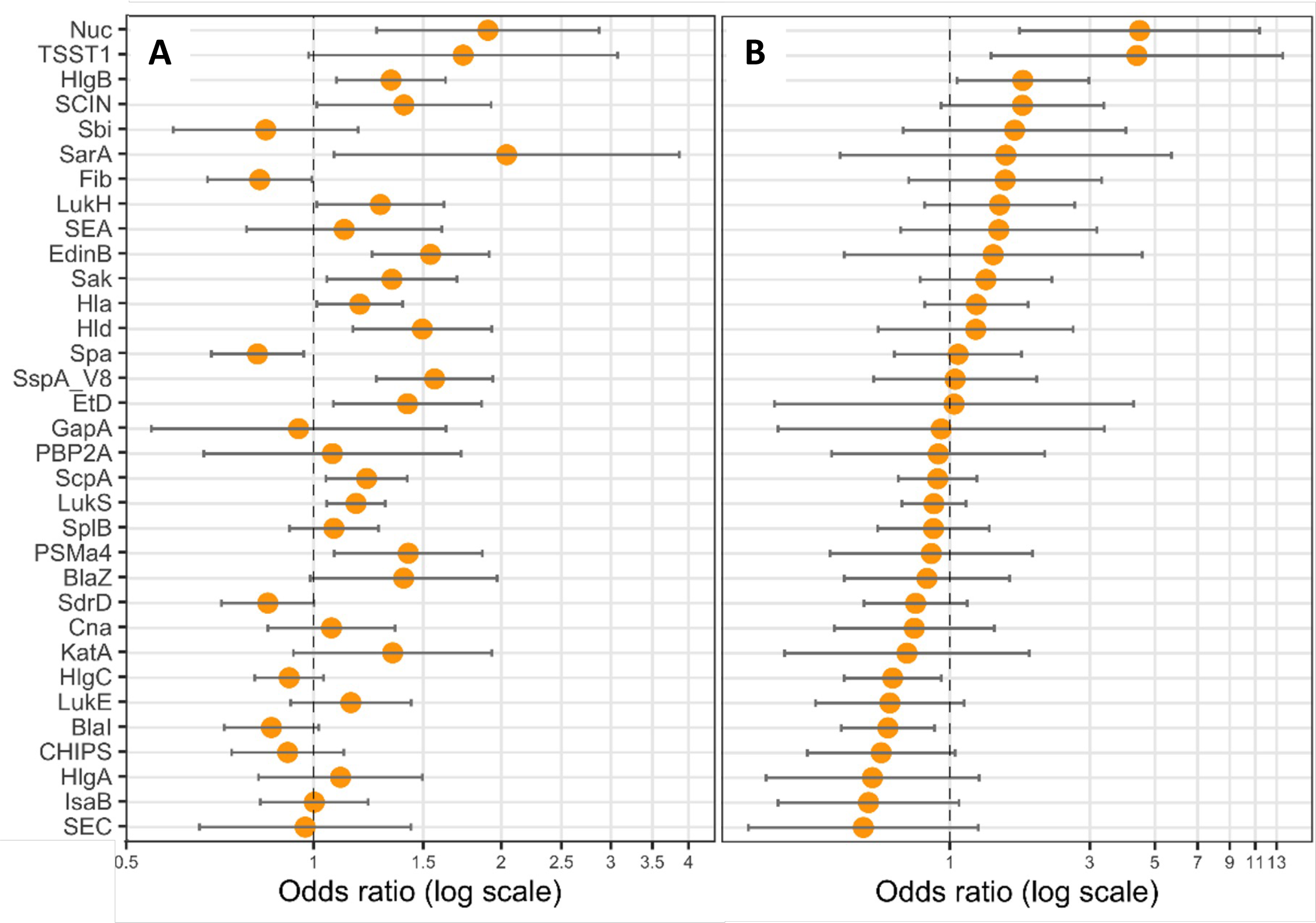
Nuc and the TSST-1 expression are correlated with leukopenia. Logistic regression models including CCI as covariate were performed, with either (**A**) one protein (univariate), or (**B**) all proteins (multivariate) as explicative variables, and leukopenia (<3G/L) as the response variable. Coefficients were reported as odds ratios (orange dots) with 95% confident intervals (error bars).

**Figure S3.**
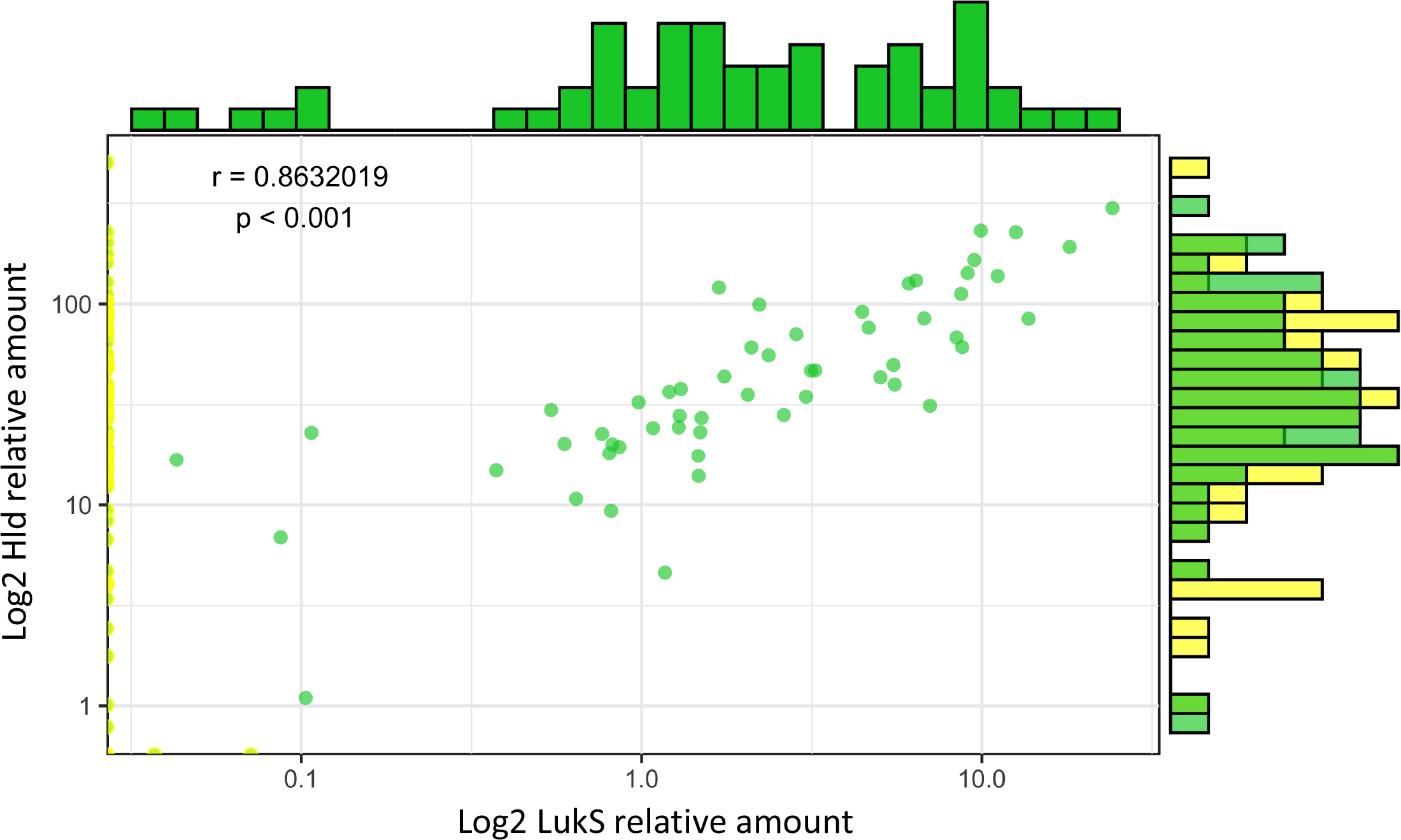
The expression levels of Hld and PVL are correlated. Log2-transformed Hla relative quantities were plotted against the log2-transformed relative quantities of LukS. The density of strains expressing similar protein quantities is represented by histograms, on top for LukS and on the right side for Hld, except for null values. Green dots represent strains with quantification for both Hld and LukS, and yellow dots represent strains with at least one null value. A Pearson correlation test was performed.

**Figure S4.**
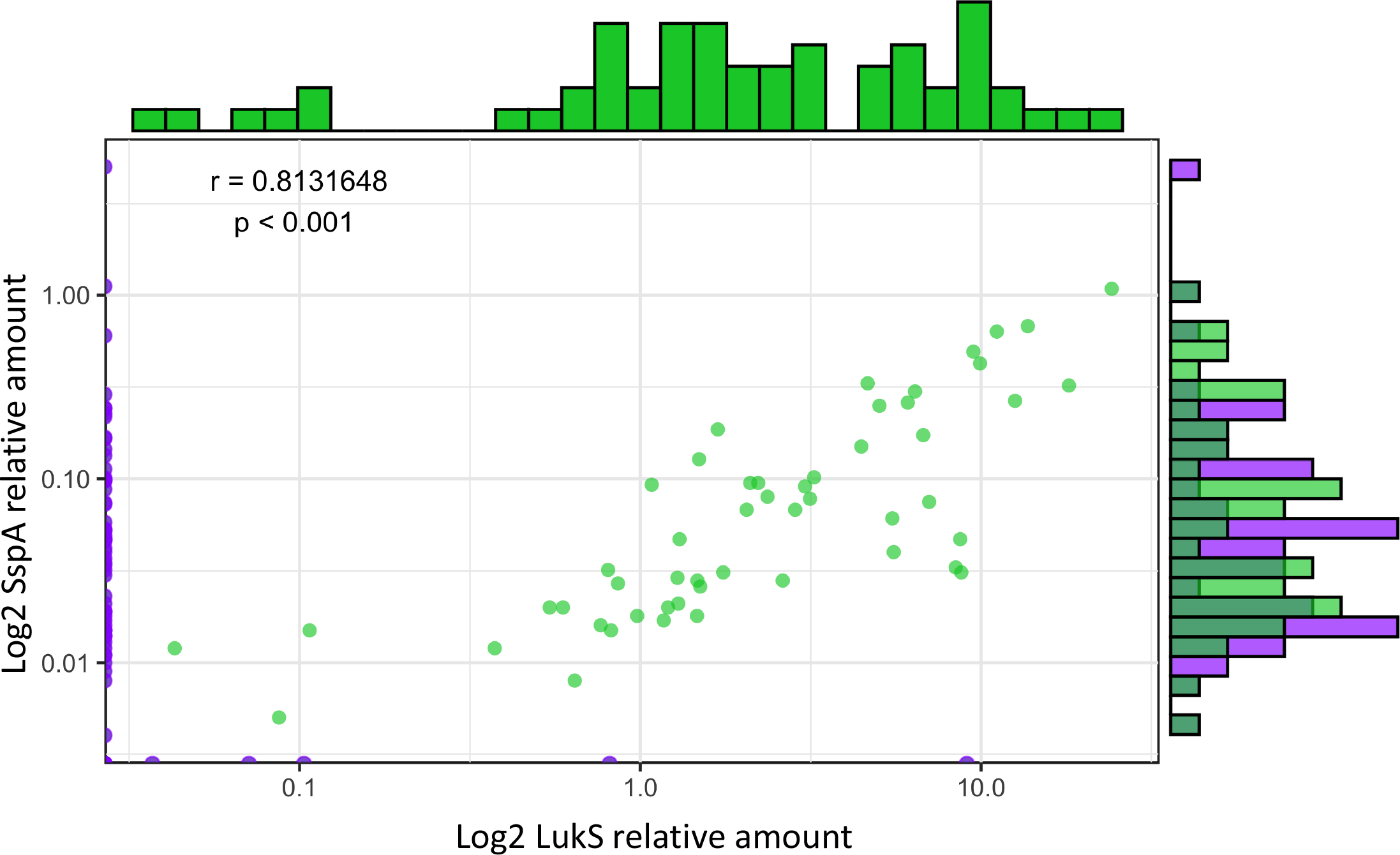
The expression levels of SspA and PVL are correlated. Log2-transformed SspA relative quantities were plotted against the log2-transformed relative quantities of LukS. The density of strains expressing similar protein quantities is represented by histograms, on top for LukS and on the right side for SspA, except for null values. Green dots represent strains with quantification for both SspA and LukS, and purple dots represent strains with at least one null value. A Pearson correlation test was performed.

Table S1 Proteins targeted and peptides used

Table S2 CA-pneumonia strains and clinical data

Table S3 Survival multivariate analysis results

Table S4 Multivariate analysis with functional groups

## Acknowledgments

We thank the scientists, engineers, and technicians of the French National Reference Center for Staphylococci for strain characterization, and Véréna Landel from the Hospices Civils de Lyon (France) for help in manuscript preparation.

## Funding

Agence Nationale de la Recherche IDAREV (JL, FV, SB)

Agence Nationale de la Recherche RHU IDBIORIV (JL, FV, SB, IM)

## Author contributions

Each author’s contribution(s) to the paper should be listed [we encourage you to follow the CRediT model]. Each CRediT role should have its own line, and there should not be any punctuation in the initials.

Conceptualization: FV, JL

Methodology: JPR, KM, FC, JL, FV

Investigation: MP, SB, IM, NM, JPR, FC

Visualization: SB, JPR

Funding acquisition: JL, FV

Project administration: JL, FV

Supervision: JL, FV

Writing – original draft: MP, IL, NM, FC, FV

Writing – review & editing: SB, JPR, KM, JL, FV

## Competing interests

Include any financial interests for each author that could be perceived as being a conflict of interest (including but not limited to financial holdings, professional affiliations, advisory positions, and board memberships). Here, also include any awarded or filed patents pertaining to the results presented in the paper, stating patent filing number and title and which authors are co-inventors. When authors have no competing interests, this should also be declared (e.g., “Authors declare that they have no competing interests.”).

FV reports research funding outside the scope of the present study by bioMérieux, two patents pending in antimicrobial resistance detection and shares in Weezion

JL reports shares in Weezion

## Data and materials availability

All data, code, and materials used in the analysis must be available in some form to any researcher for purposes of reproducing or extending the analysis. Include a note explaining any restrictions on materials, such as materials transfer agreements (MTAs). Note accession numbers to any data relating to the paper and deposited in a public database; include a brief description of the data set or model with the number. If all data are in the paper and supplementary materials, include the sentence “All data are available in the main text or the supplementary materials.”

All data are available in the main text or the supplementary materials. Whole genome sequences of the strains presented in this study will be available on ENA upon acceptance of the manuscript.

## Notes

### Competing Interest Statement

The authors have declared no competing interest.

## References

[1] Deinhardt-Emmer S, Sachse S, Geraci J, Fischer C, Kwetkat A, Dawczynski K, et al. Virulence patterns of Staphylococcus aureus strains from nasopharyngeal colonization. J Hosp Infect 2018;100:309–15. https://doi.org/10.1016/j.jhin.2017.12.011.

[2] Tong SYC, Davis JS, Eichenberger E, Holland TL, Fowler VG. Staphylococcus aureus infections: epidemiology, pathophysiology, clinical manifestations, and management. Clin Microbiol Rev 2015;28:603–61. https://doi.org/10.1128/CMR.00134-14.

[3] Foster TJ. Surface Proteins of Staphylococcus aureus. Microbiol Spectr 2019;7. https://doi.org/10.1128/microbiolspec.GPP3-0046-2018.

[4] Tam K, Torres VJ. Staphylococcus aureus Secreted Toxins and Extracellular Enzymes. Microbiol Spectr 2019;7. https://doi.org/10.1128/microbiolspec.GPP3-0039-2018.

[5] Peschel A, Otto M. Phenol-soluble modulins and staphylococcal infection. Nat Rev Microbiol 2013;11:667–73. https://doi.org/10.1038/nrmicro3110.

[6] Berger S, Kunerl A, Wasmuth S, Tierno P, Wagner K, Brügger J. Menstrual toxic shock syndrome: case report and systematic review of the literature. Lancet Infect Dis 2019;19:e313–21. https://doi.org/10.1016/S1473-3099(19)30041-6.

[7] Kulhankova K, King J, Salgado-Pabón W. Staphylococcal toxic shock syndrome: superantigen-mediated enhancement of endotoxin shock and adaptive immune suppression. Immunol Res 2014;59:182–7. https://doi.org/10.1007/s12026-014-8538-8.

[8] Brazel M, Desai A, Are A, Motaparthi K. Staphylococcal Scalded Skin Syndrome and Bullous Impetigo. Medicina (Kaunas) 2021;57:1157. https://doi.org/10.3390/medicina57111157.

[9] Recker M, Laabei M, Toleman MS, Reuter S, Saunderson RB, Blane B, et al. Clonal differences in Staphylococcus aureus bacteraemia-associated mortality. Nat Microbiol 2017;2:1381–8. https://doi.org/10.1038/s41564-017-0001-x.

[10] Deinhardt-Emmer S, Haupt KF, Garcia-Moreno M, Geraci J, Forstner C, Pletz M, et al. Staphylococcus aureus Pneumonia: Preceding Influenza Infection Paves the Way for Low-Virulent Strains. Toxins (Basel) 2019;11. https://doi.org/10.3390/toxins11120734.

[11] Bordeau V, Cady A, Revest M, Rostan O, Sassi M, Tattevin P, et al. Staphylococcus aureus Regulatory RNAs as Potential Biomarkers for Bloodstream Infections. Emerg Infect Dis 2016;22:1570–8. https://doi.org/10.3201/eid2209.151801.

[12] Stulik L, Malafa S, Hudcova J, Rouha H, Henics BZ, Craven DE, et al. α-Hemolysin Activity of Methicillin-Susceptible Staphylococcus aureus Predicts Ventilator-associated Pneumonia. Am J Respir Crit Care Med 2014;190:1139–48. https://doi.org/10.1164/rccm.201406-1012OC.

[13] Foster TJ, Geoghegan JA, Ganesh VK, Höök M. Adhesion, invasion and evasion: the many functions of the surface proteins of Staphylococcus aureus. Nat Rev Microbiol 2014;12:49–62. https://doi.org/10.1038/nrmicro3161.

[14] Carr SA, Abbatiello SE, Ackermann BL, Borchers C, Domon B, Deutsch EW, et al. Targeted Peptide Measurements in Biology and Medicine: Best Practices for Mass Spectrometry-based Assay Development Using a Fit-for-Purpose Approach *. Molecular & Cellular Proteomics 2014;13:907–17. https://doi.org/10.1074/mcp.M113.036095.

[15] Gillet Y, Tristan A, Rasigade J-P, Saadatian-Elahi M, Bouchiat C, Bes M, et al. Prognostic factors of severe community-acquired staphylococcal pneumonia in France. Eur Respir J 2021;58:2004445. https://doi.org/10.1183/13993003.04445-2020.

[16] Tran VG, Venkatasubramaniam A, Adhikari RP, Krishnan S, Wang X, Le VTM, et al. Efficacy of Active Immunization With Attenuated α-Hemolysin and Panton-Valentine Leukocidin in a Rabbit Model of Staphylococcus aureus Necrotizing Pneumonia. J Infect Dis 2020;221:267–75. https://doi.org/10.1093/infdis/jiz437.

[17] Hidalgo D, Martínez-Ortiz CA, Palsson BO, Jiménez JI, Utrilla J. Regulatory perturbations of ribosome allocation in bacteria reshape the growth proteome with a trade-off in adaptation capacity. IScience 2022;25. https://doi.org/10.1016/j.isci.2022.103879.

[18] Scott M, Klumpp Stephan, Mateescu M Eduard, Hwa Terence. Emergence of robust growth laws from optimal regulation of ribosome synthesis. Molecular Systems Biology 2014;10:747. https://doi.org/10.15252/msb.20145379.

[19] Bosdriesz E, Molenaar D, Teusink B, Bruggeman FJ. How fast-growing bacteria robustly tune their ribosome concentration to approximate growth-rate maximization. The FEBS Journal 2015;282:2029–44. https://doi.org/10.1111/febs.13258.

[20] Cecchini T, Yoon E-J, Charretier Y, Bardet C, Beaulieu C, Lacoux X, et al. Deciphering Multifactorial Resistance Phenotypes in Acinetobacter baumannii by Genomics and Targeted Label-free Proteomics *. Molecular & Cellular Proteomics 2018;17:442–56. https://doi.org/10.1074/mcp.RA117.000107.

[21] Laabei M, Peacock SJ, Blane B, Baines SL, Howden BP, Stinear TP, et al. Significant variability exists in the cytotoxicity of global methicillin-resistant Staphylococcus aureus lineages. Microbiology (Reading) 2021;167. https://doi.org/10.1099/mic.0.001119.

[22] Bronesky D, Wu Z, Marzi S, Walter P, Geissmann T, Moreau K, et al. Staphylococcus aureus RNAIII and Its Regulon Link Quorum Sensing, Stress Responses, Metabolic Adaptation, and Regulation of Virulence Gene Expression. Annual Review of Microbiology 2016;70:299–316. https://doi.org/10.1146/annurev-micro-102215-095708.

[23] Felden B, Bouloc P. Regulatory RNAs in bacteria: From identification to function. Methods 2017;117:1–2. https://doi.org/10.1016/j.ymeth.2017.03.018.

[24] Fechter P, Caldelari I, Lioliou E, Romby P. Novel aspects of RNA regulation in Staphylococcus aureus. FEBS Lett 2014;588:2523–9. https://doi.org/10.1016/j.febslet.2014.05.037.

[25] Bronner S, Monteil H, Prévost G. Regulation of virulence determinants in Staphylococcus aureus: complexity and applications. FEMS Microbiology Reviews 2004;28:183–200. https://doi.org/10.1016/j.femsre.2003.09.003.

[26] Geisinger E, Adhikari RP, Jin R, Ross HF, Novick RP. Inhibition of rot translation by RNAIII, a key feature of agr function. Mol Microbiol 2006;61:1038–48. https://doi.org/10.1111/j.1365-2958.2006.05292.x.

[27] Reyes D, Andrey DO, Monod A, Kelley WL, Zhang G, Cheung AL. Coordinated regulation by AgrA, SarA, and SarR to control agr expression in Staphylococcus aureus. J Bacteriol 2011;193:6020–31. https://doi.org/10.1128/JB.05436-11.

[28] Xue T, You Y, Shang F, Sun B. Rot and Agr system modulate fibrinogen-binding ability mainly by regulating clfB expression in Staphylococcus aureus NCTC8325. Med Microbiol Immunol 2012;201:81–92. https://doi.org/10.1007/s00430-011-0208-z.

[29] Zhou Y, Niu C, Ma B, Xue X, Li Z, Chen Z, et al. Inhibiting PSMα-induced neutrophil necroptosis protects mice with MRSA pneumonia by blocking the agr system. Cell Death Dis 2018;9:1–14. https://doi.org/10.1038/s41419-018-0398-z.

[30] Alonzo F, Torres VJ. The Bicomponent Pore-Forming Leucocidins of Staphylococcus aureus. Microbiol Mol Biol Rev 2014;78:199–230. https://doi.org/10.1128/MMBR.00055-13.

[31] Gillet Y, Issartel B, Vanhems P, Fournet J-C, Lina G, Bes M, et al. Association between Staphylococcus aureus strains carrying gene for Panton-Valentine leukocidin and highly lethal necrotising pneumonia in young immunocompetent patients. Lancet 2002;359:753–9. https://doi.org/10.1016/S0140-6736(02)07877-7.

[32] Lina G, Piémont Y, Godail-Gamot F, Bes M, Peter M-O, Gauduchon V, et al. Involvement of Panton-Valentine Leukocidin—Producing Staphylococcus aureus in Primary Skin Infections and Pneumonia. Clin Infect Dis 1999;29:1128–32. https://doi.org/10.1086/313461.

[33] Spaan AN, Henry T, van Rooijen WJM, Perret M, Badiou C, Aerts PC, et al. The Staphylococcal Toxin Panton-Valentine Leukocidin Targets Human C5a Receptors. Cell Host & Microbe 2013;13:584–94. https://doi.org/10.1016/j.chom.2013.04.006.

[34] Diep BA, Chan L, Tattevin P, Kajikawa O, Martin TR, Basuino L, et al. Polymorphonuclear leukocytes mediate Staphylococcus aureus Panton-Valentine leukocidin-induced lung inflammation and injury. PNAS 2010;107:5587–92. https://doi.org/10.1073/pnas.0912403107.

[35] Pivard M, Moreau K, Vandenesch F. Staphylococcus aureus Arsenal To Conquer the Lower Respiratory Tract. MSphere 2021;6:e00059–21. https://doi.org/10.1128/mSphere.00059-21.

[36] Spaan AN, Vrieling M, Wallet P, Badiou C, Reyes-Robles T, Ohneck EA, et al. The staphylococcal toxins γ-haemolysin AB and CB differentially target phagocytes by employing specific chemokine receptors. Nature Communications 2014;5:5438. https://doi.org/10.1038/ncomms6438.

[37] Spaan AN, Schiepers A, Haas CJC de, Hooijdonk DDJJ van, Badiou C, Contamin H, et al. Differential Interaction of the Staphylococcal Toxins Panton–Valentine Leukocidin and γ-Hemolysin CB with Human C5a Receptors. The Journal of Immunology 2015;195:1034–43. https://doi.org/10.4049/jimmunol.1500604.

[38] Thammavongsa V, Missiakas DM, Schneewind O. Staphylococcus aureus degrades neutrophil extracellular traps to promote immune cell death. Science 2013;342:863–6. https://doi.org/10.1126/science.1242255.

[39] Waclavicek M, Stich N, Rappan I, Bergmeister H, Eibl MM. Analysis of the early response to TSST-1 reveals Vbeta-unrestricted extravasation, compartmentalization of the response, and unresponsiveness but not anergy to TSST-1. J Leukoc Biol 2009;85:44–54. https://doi.org/10.1189/jlb.0108074.

[40] Hackbarth CJ, Chambers HF. blaI and blaR1 regulate beta-lactamase and PBP 2a production in methicillin-resistant Staphylococcus aureus. Antimicrobial Agents and Chemotherapy 1993;37:1144–9. https://doi.org/10.1128/AAC.37.5.1144.

[41] Croes S, Beisser PS, Terporten PH, Neef C, Deurenberg RH, Stobberingh EE. Diminished in vitro antibacterial activity of oxacillin against clinical isolates of borderline oxacillin-resistant Staphylococcus aureus. Clin Microbiol Infect 2010;16:979–85. https://doi.org/10.1111/j.1469-0691.2010.02956.x.

[42] Hryniewicz MM, Garbacz K 2017. Borderline oxacillin-resistant Staphylococcus aureus (BORSA) – a more common problem than expected? Journal of Medical Microbiology n.d.;66:1367–73. https://doi.org/10.1099/jmm.0.000585.

[43] Rouha H, Badarau A, Visram ZC, Battles MB, Prinz B, Magyarics Z, et al. Five birds, one stone: neutralization of α-hemolysin and 4 bi-component leukocidins of Staphylococcus aureus with a single human monoclonal antibody. MAbs 2015;7:243–54. https://doi.org/10.4161/19420862.2014.985132.

[44] Ludwig C, Claassen M, Schmidt A, Aebersold R. Estimation of absolute protein quantities of unlabeled samples by selected reaction monitoring mass spectrometry. Mol Cell Proteomics 2012;11:M111.013987. https://doi.org/10.1074/mcp.M111.013987.

[45] Rasigade J-P, Barray A, Shapiro JT, Coquisart C, Vigouroux Y, Bal A, et al. A viral perspective on worldwide non-pharmaceutical interventions against COVID-19 2020:2020.08.24.20180927. https://doi.org/10.1101/2020.08.24.20180927.

